# Modulating Brain Perfusion, Functional Connectivity, and Metabolite Patterns through Theta Burst Transcranial Ultrasonic Stimulation

**DOI:** 10.1101/2025.06.25.661571

**Authors:** Daniel Keeser, Lukas Roell, Veronica Meedt, Maximilian Haßlberger, Maxim Korman, Enrico Schulz, Maximilian Lueckel, Berkhan Karsli, Genc Hasanaj, Theresa Faeßler, Gizem Vural, Kai-Yen Chang, Lucia Bulubas, Frank Padberg, Florian Raabe, Peter Falkai, Til Ole Bergmann, Boris-Stephan Rauchmann

## Abstract

**Background:** Transcranial ultrasonic stimulation (TUS) is an emerging non-invasive neuromodulation technique with the potential to target both cortical and subcortical brain regions. This study investigates the effects of theta-burst TUS (tb-TUS), a neuromodulatory pattern characterized by bursts of pulses repeated at a theta frequency, on cerebral blood flow, functional connectivity, and metabolite concentrations in the primary motor cortex (M1). The aim of this study is to take a first step towards the mechanistic and methodological feasibility of tb-TUS at the M1 using multimodal neuroimaging.

**Methods:** Seventeen healthy participants underwent a double-blind, sham-controlled crossover design, receiving both active and sham tb-TUS to the left M1 over three days. Multimodal MRI, including pseudo-continuous arterial spin labeling (PCASL), resting-state functional MRI (rs-fMRI), and magnetic resonance spectroscopy (MRS), was conducted at baseline, pre-, and post-stimulation. Acoustic simulations and finger-tapping BOLD-peak signal guided individualized TUS targeting.

**Results:** Active tb-TUS significantly reduced cerebral blood flow (p < .001) and within-region functional connectivity (p < .001) in the M1 compared to sham stimulation. A non-significant trend towards decreased GABA was observed, with no significant session x condition interaction found for GABA, Glutamate, or Glx concentrations.

**Conclusion:** This pilot study demonstrates that tb-TUS of the M1 induces reductions in cerebral blood flow and functional connectivity in healthy participants. Our findings indicate that tb-TUS may be mitigating neural hyperactivity patterns, but preliminary studies so far arrive at differing results, highlighting the need for further research to replicate our findings, elucidate the underlying mechanisms, and optimize stimulation protocols.

## Introduction

### Introduction to Transcranial Ultrasonic Stimulation (TUS)

Among currently available non-invasive brain stimulation techniques, transcranial ultrasonic stimulation (TUS) has gained growing interest in recent years [1,2]. TUS is unique in its ability to non-invasively target both cortical and subcortical brain regions with high spatial precision. This capability largely confines the intended neuromodulatory effects to the focal area, while minimizing substantial direct impact on adjacent non-targeted tissue [3–6]. As revealed by multiple examinations conducted in animals and humans, TUS can be considered safe with only minor side effects reported [7]. Consequently, recent studies have investigated the ability of TUS to modulate brain functioning in both animals and humans, aiming to explore its potential future therapeutic use in psychiatric and neurological patients [8]. Initial clinical TUS pilot studies have yielded promising, albeit preliminary, results across several domains [9].

### Targeting Dysfunctional Brain Networks

Transcranial focused ultrasonic stimulation (TUS), which enables the precise modulation of deep brain circuits, shows therapeutic potential for several challenging disorders. Recent work highlights that symptom relief is often achieved by targeting key hubs within dysfunctional brain networks. This is well-illustrated in major depression studies, where targeting nodes within the brain’s mood and default-mode networks has successfully reduced symptoms. For example, a randomized trial demonstrated that modulating the subcallosal cingulate cortex (SCC) improved depression scores [10]. Another highly targeted trial, also focusing on the SCC in treatment-resistant patients, reported a significant reduction in symptoms [11]. This circuit-based approach is also being explored for other conditions. Initial studies have shown cognitive gains in Alzheimer’s disease after targeting the hippocampus [12], as well as symptom alleviation in chronic pain and schizophrenia through the modulation of specific cortical regions [13,14]. While these pioneering studies highlight the therapeutic potential of TUS, it is crucial to recognize their preliminary nature.

Deploying TUS on different neural targets in animals has indicated that it can elicit excitatory and inhibitory neuronal responses depending on the target regions stimulated and the stimulation protocol applied [15,16]. This has been confirmed in several human trials, demonstrating that TUS on the one hand may lead to excitatory activation patterns in the motor-somatosensory cortex as derived from electroencephalography (EEG) in healthy subjects [17], but can also induce an inhibition of thalamic nuclei, assessed through somatosensory evoked potentials and accompanied by worsened performance in discrimination tasks [4,5]. Therefore, further studies confirming the differential effects of stimulation protocols are urgently needed. Here, we employed “theta-burst stimulation”, a neuromodulatory pattern characterized by bursts of pulses repeated at a theta frequency, typically around 5 Hz. Although recent work suggests that ’5Hz repetitive TUS (5Hz-rTUS)’ is a more specific descriptor for this protocol [18], we use the term “theta-burst” for consistency. Our terminology aligns with its use in numerous foundational studies to which we refer (e.g. [19–21]).

In a recent study by Zeng et al., theta burst TUS (tb-TUS) to the left primary motor cortex (M1) effectively induced corticospinal excitability and changes in intracortical inhibition and facilitation, whereas regular TUS and sham tb-TUS did not show these effects [19].

Our study builds on this preliminary work by examining the effects of tb-TUS to the left M1 on cerebral blood flow, resting-state functional MRI connectivity, and metabolite concentrations.

### Rationale for this study

To leverage future large-scale studies in this field, further basic research on the mechanistic effects of TUS in the healthy brain is warranted. While the reviewed clinical pilot trials across various brain regions and patient populations are encouraging, they also highlight the preliminary nature of current TUS applications and the need for a more systematic understanding of its neuromodulatory effects. The variability in targets, patient characteristics, and stimulation protocols underscores the importance of basic research to better understand the precise physiological impact of TUS, particularly for novel paradigms like theta-burst stimulation. To contribute to such an evidence base, it is crucial to first characterize these effects in a well-controlled manner in the healthy human brain and in a well-understood brain area that is subject to less variability, such as the frontal brain. Therefore, we conducted a comprehensive repeated measures design comparing TUS of the motor cortex with sham stimulation in healthy subjects. Our aim was to investigate the effects of TUS on cerebral blood flow, functional connectivity and neurotransmitter concentrations in the motor cortex, respectively.

## Methods

The present study was conducted at the Department of Psychiatry and Psychotherapy of the Ludwig-Maximilians-University (University Hospital LMU) in Munich in accordance with the Declaration of Helsinki. Ethical approval was provided by the local ethics committee of LMU Munich (23-0116).

### Study Sample

17 healthy subjects (nine females) aged 22–39 years (mean = 27.06, SD = 4.47) were included in the current study. Individuals under legal guardianship or with limited ability to consent were excluded. Additional exclusion criteria encompassed psychiatric disorders (e.g., schizophrenia, bipolar disorder, major depression), past or current substance abuse, and acute suicidal behavior. Participants with a history of severe traumatic brain injury, structural damage to the basal ganglia or brainstem, neurological conditions (e.g., epilepsy, cerebrovascular disease), or severe internal diseases (e.g., cardiac, hepatic, or respiratory conditions) were also excluded. Other criteria included severe infectious, systemic skin, or bone diseases (e.g., osteoporosis), as well as malignant brain conditions such as tumors. Due to MRI constraints, individuals with electronic implants (e.g., pacemakers) or other contraindications were not eligible. All participants provided written informed consent. For this pilot study we used a sample size of N=17 participants. This number is consistent with sample sizes commonly employed in exploratory neuroimaging research investigating the acute effects of novel non-invasive brain stimulation techniques, including transcranial ultrasonic stimulation (TUS) studies that utilize multimodal imaging approaches. Such a sample size was deemed adequate for the primary aims of this study: to detect initial, potentially medium-to-large, neurophysiological effects of theta-burst TUS on cerebral blood flow, functional connectivity, and metabolite concentrations, and to provide preliminary data for effect size estimation to inform the design and power calculations of future, larger-scale investigations.

### Study Design

This study followed a double-blind, sham-controlled crossover design. Each participant received both sham and active theta-burst TUS stimulation in a randomized order over the course of three days. The sequence of these conditions (active first or sham first) was determined for each participant using computer-generated random numbers. This randomization procedure assigned the order of conditions without any further restrictions, such as blocking. MRI scans were conducted at baseline, immediately before, and after stimulation, following a standardized protocol at the same time each day (±30 min) to control for circadian effects. At baseline, each participant underwent a finger-tapping task (Figure 1), and the individual maximum activation at the left M1, determined from the task-related fMRI, was used to guide the simulation and active stimulation targeting (see Figure S2).

**Figure 1.**
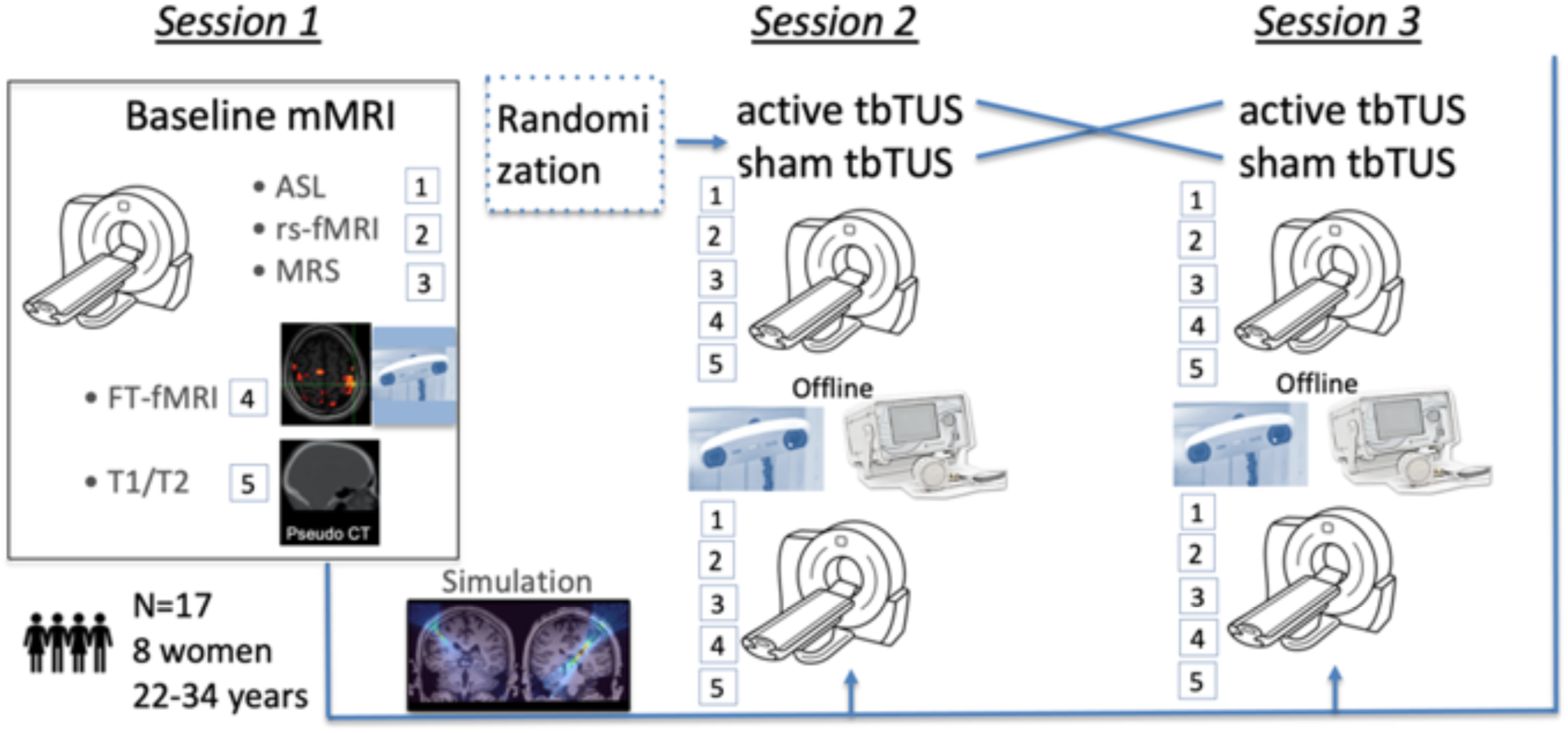
Experimental Design. Seventeen healthy volunteers (mean age=26.31, SD=3.34; 8 women). Repeated measurement design over three days, including baseline and randomized active/sham theta burst TUS (tbTUS) stimulation conditions in a cross-over design. Multimodal MRI imaging including 1. pcASL, 2. rs-fMRI, 3.MRS, 4.FT-fMRI and structural 5.T1w/T2w scans. Simulations were performed using k-wave toolbox [31].

In the active condition the left primary motor cortex was stimulated, while sham stimulation was applied to the left ventricular space, as suggested recently as a proper sham condition in TUS studies [22]. TUS targeting was individualized using T1-weighted MRI scans. The study design is illustrated in Figure 1.

### Theta-Burst TUS Stimulation Guided by Neuronavigation

Participants were seated semi-upright in a reclining chair with head stabilization using vacuum padding (see Figure S1). Theta-burst stimulation was delivered using the four-element NeuroFUS PRO - CTX-500 transducer (500 kHz; Brainbox Ltd., Cardiff, UK). To ensure precise positioning, the transducer was guided by a neuronavigation system (LOCALITE, Bonn, Germany) based on each participant’s T1-weighted MRI. Before stimulation ultrasound gel was applied for optimal transmission. The experimenter held the transducer in place, with real-time neuronavigation feedback ensuring accurate targeting of the motor cortex or ventricle. For both sham and active TUS stimulation, the transducer remained in the same position and orientation to avoid differential auditory confounds. To ensure blinding for each specific stimulation session, an unblinded member of the research team (who was not involved in outcome assessment or participant interaction during data acquisition) was responsible for positioning the transducer. This individual would set the device to deliver either active or sham stimulation according to the participant’s allocated sequence for that particular day. The transducer and setup were identical for both conditions so that the active condition could not be distinguished from the sham condition. The experimenter delivering the TUS stimulation, the participants, and all personnel involved in data acquisition and analysis remained blind to the condition assigned for each session throughout the study. The sonication parameters are shown in Table S1.

### Acoustic Simulation

Before TUS stimulation, spatial targeting was guided by T1-weighted MRI-based acoustic simulations to ensure sufficient energy delivery while maintaining safety standards. This precise targeting is crucial due to the skull’s heterogeneities in thickness and speed of sound. Simulations were performed individually using the k-Wave toolbox [23] in MATLAB, with pseudo-CT scans generated via the MR-to-pCT for FUS acoustic simulations code [32] to carry out acoustic simulations based on individual shape and acoustic properties of the skull. The *BRIC_TUS_Simulation_Tools* framework [24] was used to integrate transducer geometry, ultrasonic beam visualization, and transducer phase parameters for different focus depths. To match the source pressure amplitude of the transducer between k-Wave simulations and the device, a lookup table of ISPPA-source pressure pairs was created, using water tank simulations to determine ISPPA values at the focal target. The transducer amplitude was then set to match the respective ISPPA obtained during prior scan tank measurements by the device manufacturer. Focus coordinates were visualized in MATLAB R2023a (MathWorks, Natick, MA, USA), with an orthogonal axis manually set to align the focus point and transducer. Due to the restricted minimal focus depth (34 mm) of the hardware, an offset was applied to ensure proper targeting, which was realized using ultrasonic gel pads (Parker, Fairfield, NJ, USA) to match the additional offset in simulations.

For sham stimulation, gel pad thicknesses were chosen similarly as in the active setup, where the device’s focus depth needed to be maximized (67 mm) for each subject, while requiring an offset between transducer and skull slightly below that of the active condition to prevent cortical gray matter stimulation (> -6dB). The transducer axis remained consistent between conditions. Acoustic simulations were iteratively refined until the active tb-TUS intracranial spatial peak temporal average intensity (ISPTA), a common measure of how much ultrasound energy is being delivered to the brain tissue, fell within 620–720 mW/cm². Sham stimulation used the same source pressure amplitudes, yielding significantly lower ISPTA as a consequence of the elongated focal beam and larger focus depth. Visualized acoustic simulations for both conditions are shown in Figure 2.

**Figure 2.**
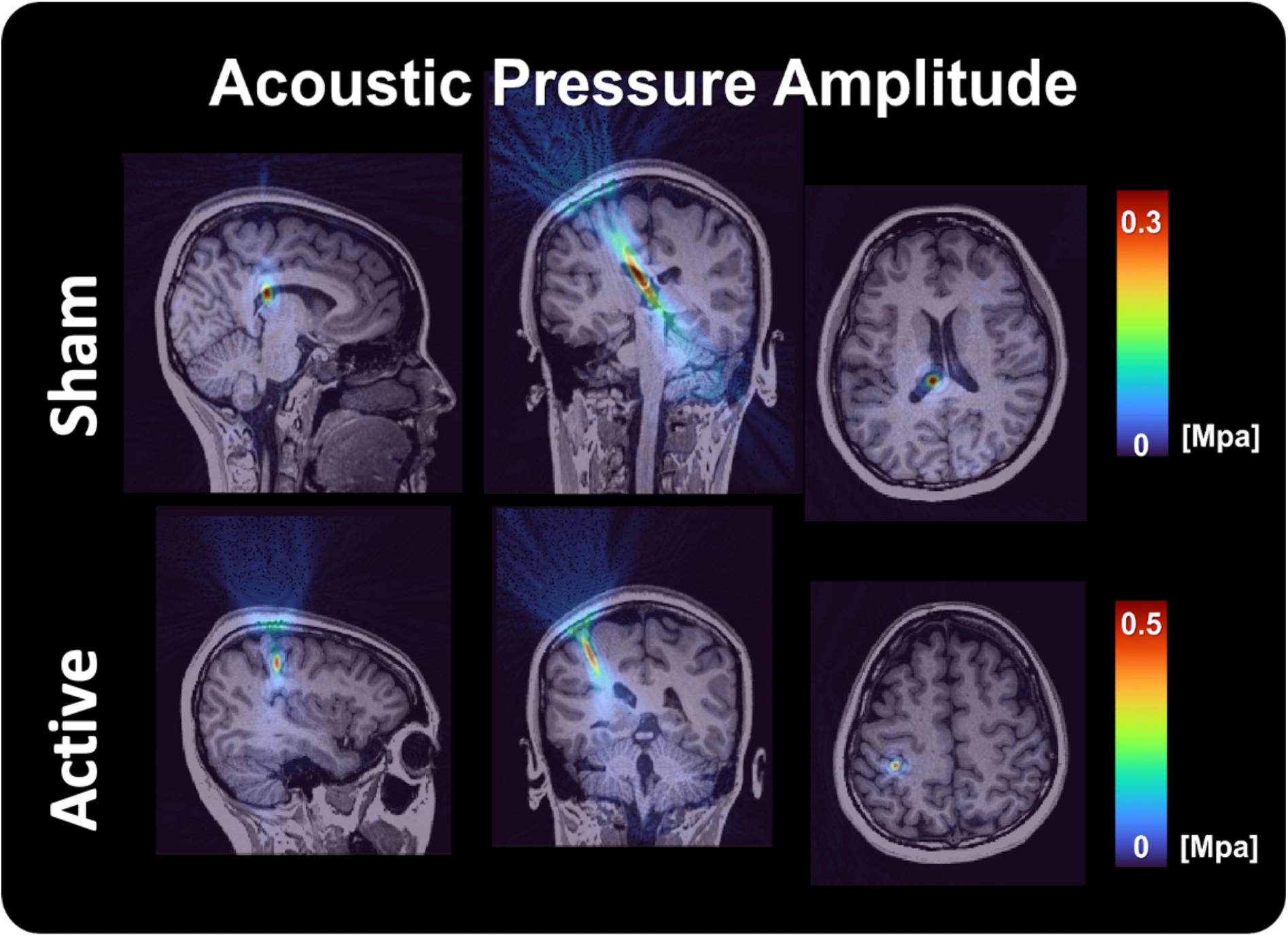
Acoustic sham simulation and active theta-burst TUS stimulation. A simulated ultrasound pressure field superimposed on a T1-weighted MRI shows precise targeting of the left ventricle for sham stimulation (top row) and the motor cortex (bottom row) for active stimulation. Simulations were done with k-wave [23].

### Multimodal MRI Data Acquisition and Processing

MRI was performed at the Neuro-Imaging-Core-Unit-Munich (NICUM) at the Department of Psychiatry and Psychotherapy of the LMU Hospital Munich using a 3T Siemens Magnetom Prisma scanner (Siemens Healthcare, Erlangen) with a 32-channel head coil. Each participant underwent five MRI sessions each including T1w MRI, PCASL, rs-fMRI, and MRS. A detailed description of scanning protocols summarized in Table S2.

PCASL data was processed using Oxford ASL resulting in measures of regional cerebral blood flow quantified in milliliters per 100g brain tissue per minute [25,26]. Twenty-four brain regions of interest within the motor cortex were selected from the Brainnetome Atlas [27,28], focusing on the Precentral Gyrus (PrG), Postcentral Gyrus (PoG), and Paracentral Lobule (PCL). These regions are visualized in Figure 3A. We utilized the mean cerebral blood flow across these regions as primary outcome. With regard to the MRS data, a CMRR msLaser sequence was deployed with a 20×20×20 mm voxel positioned in the primary motor cortex, based on [29]. MRS analysis was performed in MATLAB using the Osprey (v2.5.0) [30] and LCModel (Linear Combination Model, v6.3-1R) [31]. Glx and GABA concentrations were used as final outcomes from MRS.

**Figure 3.**
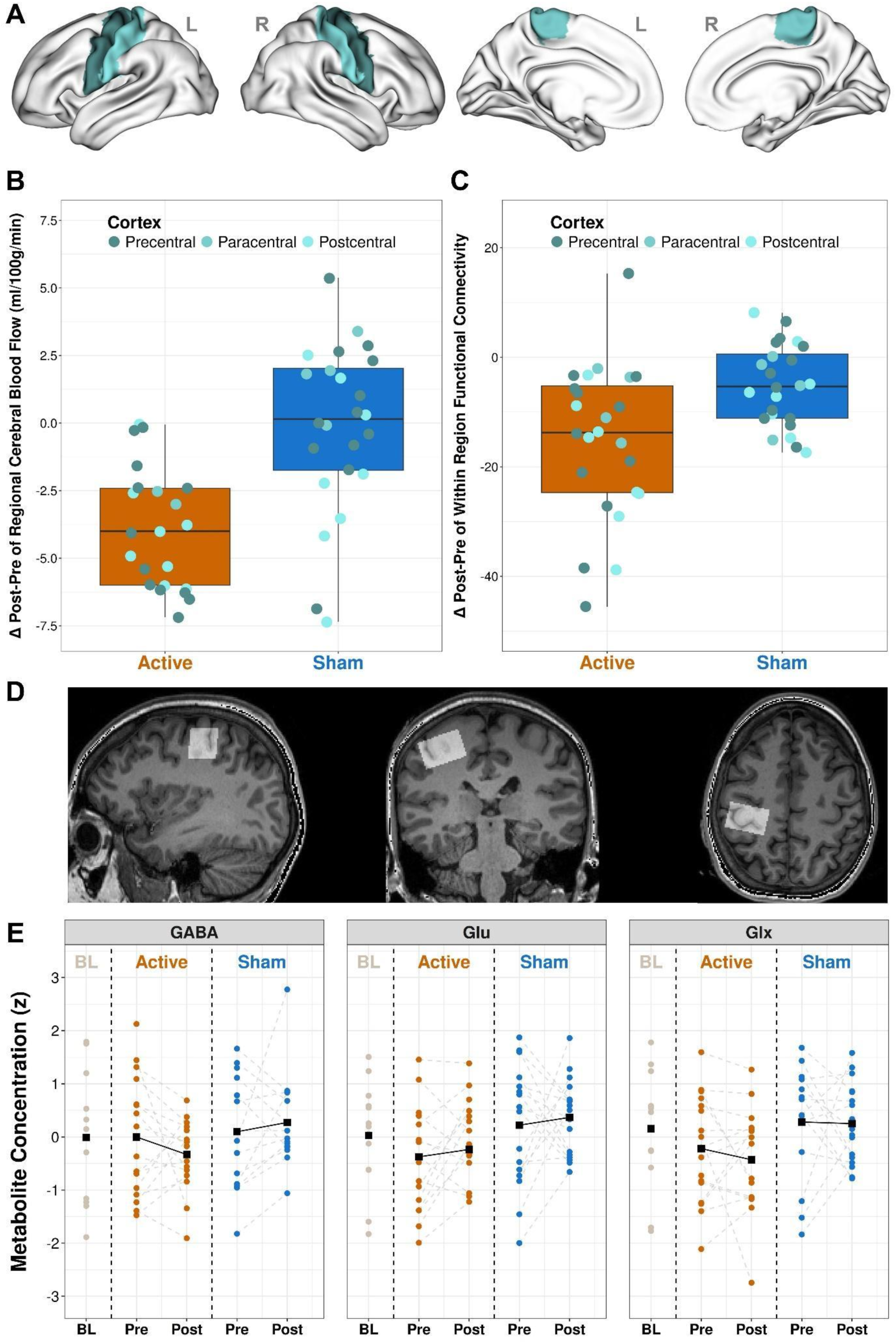
Effects of theta-burst TUS stimulation on cerebral blood flow, functional MRI connectivity, and metabolite concentrations. (A) Surface views of the left and right somato-motor cortex (precentral, paracentral and postcentral Brainnetome [35] ROIs) that were used for data analysis. (B) Magnitude of Changes between pre and post stimulation in Cerebral Blood flow following active stimulation compared to sham. Bar graph quantifying the mean change (post-stimulation minus pre-stimulation) in rCBF (ml/100g/min) extracted from significantly altered regions, comparing active (orange) and sham (blue) conditions. (C) Bar graph showing the mean change (post-stimulation minus pre-stimulation) in within-region functional connectivity for the precentral, paracentral, and postcentral gyri, comparing active (orange) and sham (blue) stimulation. (D) Localisation of the MRS ROI (20x30x20mm) on the left primary Cortex (M1) shown on an example subject’s T1w-MPRAGE. (E) Individual participant changes in metabolite concentrations. Plots display concentrations for Gamma-aminobutyric acid (GABA), Glutamate (Glu), and the composite of Glutamate+Glutamine (Glx). For both active and sham conditions, each line connects a single subject’s pre- and post-stimulation measurements, with baseline (BL) shown for reference. Concentrations are represented as z-scores relative to baseline.

Processing of the rs-fMRI data was conducted using FSL [32] and the Neuromodulation and Multimodal Neuroimaging Software (NAMNIs) [28]. Functional connectivity was calculated within the aforementioned regions of the motor cortex from the Brainnetome Atlas [27].

Details on the analysis steps and quality control procedures applied in each modality are described in the supplemental information, Table S3.

### Statistical Data Analysis

Statistical analysis was performed in R version 4.4.1 [33]. To examine the effect of TUS on the cerebral blood flow in the motor cortex, we computed a linear mixed effect model for repeated measures using the *lme4* package [34]. We included session (pre vs. post stimulation), condition (active vs. sham stimulation), the session x condition interaction, age, sex, and motor cortex region (24 regions defined by the Brainnetome atlas [27]) as fixed effects, while the cerebral blood flow served as dependent variable. A random intercept was modelled for each subject. In the case of a significant session x condition interaction (p < 0.05), Tukey post-hoc tests were calculated to identify the direction and size of the effect from pre to post intervention for each condition.

The same statistical approach was applied to within-region functional connectivity as dependent variable derived from rs-fMRI to investigate the impact of TUS on motor cortex functional connectivity.

For MRS, we computed three separate linear mixed effect models with GABA, Glutamate, and Glx concentrations in the motor cortex as dependent variables and session, condition, the session x condition interaction, age, and sex as fixed effects. A random intercept was again modelled for each subject. Tukey post-hoc tests were again calculated if the significant session x condition interaction was significant (p < 0.05).

## Results

The main findings are visualized in Figure 3. Finger tapping peak activations of three random subjects of the sample are shown in Figure S2. Additional figures of our results are provided in the supplemental information.

### Effects of TUS on Cerebral Blood Flow in the Motor Cortex

We obtained a significant session x condition interaction (F = 5.96, p = 0.015), suggesting a difference between active and sham stimulation regarding the change of the cerebral blood flow of the motor cortex from pre to post session. Tukey post-hoc tests revealed a reduction of cerebral blood flow in the motor cortex after active stimulation (d = -0.25, CI = [-0.39, -0.12], p < 0.001), but not after sham stimulation (d = -0.01, CI = [-0.15, 0.13], p = 0.877).

### Effects of TUS on Functional Connectivity within the Motor Cortex

Building on the finding that tb-TUS altered local cerebral blood flow, we next aimed to understand if this local modulation also influenced functional connectivity within the stimulated region. A significant session x condition interaction was found (F = 3.88, p = 0.049), indicating a distinct pattern of functional connectivity change within the motor cortex from pre to post session between active and sham stimulation. Tukey post-hoc tests demonstrated a reduction of within-region functional connectivity in the motor cortex after active stimulation (d = -0.30, CI = [-0.43, -0.16], p < 0.001), but not after sham stimulation (d = -0.10, CI = [-0.24, 0.04], p = 0.145).

### Effects of TUS on GABA, Glutamate, and Glx Concentrations in the Motor Cortex

After observing significant changes at both the vascular and network levels, we proceeded to investigate the potential underlying neurochemical mechanisms. Despite a descriptive tendency for GABA (Figure 3), no significant session x condition interaction was obtained for GABA (F = 2.08, p = 0.157), Glutamate (F = 0.00, p = 0.984), or Glx (F = 0.04, p = 0.838), suggesting no difference in concentration change between active and sham stimulation.

### Safety and Tolerability

A final and critical component of our study was to ensure the safety of the TUS application. To provide an objective assessment of any potential TUS-induced brain injury, an experienced neuroradiologist, blinded to the stimulation condition, systematically reviewed post-stimulation structural MRI scans. The imaging protocol included T1-weighted and T2-weighted sequences, FLAIR sequences to evaluate for edema, and specific sequences sensitive to microhemorrhages (i.e., T2*-weighted gradient echo or Susceptibility-Weighted Imaging (SWI)). Our neuroradiological evaluation revealed no evidence of edema, hemorrhage, or any other acute structural changes attributable to the TUS application in any participant.

## Discussion

This pilot study provides preliminary evidence that transcranial ultrasonic stimulation with a theta-bursts protocol (tb-TUS) in the primary motor cortex (M1) leads to quantifiable neurophysiological changes. Our principal findings were a significant reduction in local cerebral blood flow (CBF), measured by pseudo-continuous arterial spin labeling (PCASL), and a concurrent decrease in within-region functional connectivity (WR-FC), measured by resting-state fMRI. In addition to these hemodynamic changes, magnetic resonance spectroscopy (MRS) showed a trend toward reduced GABA concentrations, although this effect was not statistically significant. Our finding is consistent with Yaakub et al. [21] but stands in contrast to the significant inhibitory effects found in the other two modalities.

The observed decreases in CBF and local FC are physiologically consistent. This relationship is likely bidirectional: while a reduction in regional FC (and thus neural activity) is expected to decrease local blood flow via neurovascular coupling, it is also plausible that a primary reduction in CBF could disrupt the metabolic environment necessary for neuronal synchrony, thereby causing a secondary decrease in FC. Our findings thus provide a coherent set of initial neuroimaging markers reflecting a net suppressive effect of this tb-TUS protocol on local cortical function in M1. These findings are consistent with recent human studies, such as the work by Cain et al., who also reported a significant reduction in both BOLD signal and ASL-measured perfusion in deep-brain structures following low-intensity transcranial ultrasonic stimulation [35]. This lends further support to the view that this modality can induce a net inhibitory effect on local brain activity.

Further indirect support for our findings comes from electrophysiological data. Darmani et al. recently showed that tb-TUS applied to the globus pallidus specifically enhances power of local field potentials in the theta EEG band [36] of Parkinson and dystonia patients. This is a crucial observation, as an increase in theta oscillations represents a slowing of neural activity, which has been independently associated with reductions in the BOLD signal [37]. The combination of electrophysiological and hemodynamic findings in first pilot studies may be interpreted as evidence for an inhibitory effect of tb-TUS on local neural activity.

It is critical, however, to place these findings within the complex and evolving landscape of tb-TUS research. The suppressive effects we observed on hemodynamic and connectivity measures contrast with other studies using similar tb-TUS protocols on M1 that reported *increased* corticospinal excitability, measured by motor-evoked potentials (MEPs) [19,20], whereas Bao et al. found reduced MEPs with a continuous tb-TUS protocol and no change with intermittent tb-TUS stimulation [38]. Accordingly, the temporal pattern of stimulation might be a contributing factor, as Kim et al. recently demonstrated that while an *intermittent* tb-TUS protocol led to potentiation, a *continuous* protocol produced a lasting suppression on MEPs in mice [39]. This divergence underscores that different measurement modalities capture distinct aspects of the neural response (see Table S4 for an overview of tb-TUS studies with similar protocols on MEPs and imaging modalities). The variability is not just between modalities and different studies but can also be seen within a single study. For instance, Barksdale et al., while reporting a significant *group-level* reduction in amygdala BOLD signal with TUS, also noted considerable inter-individual variability, with a subset of participants showing an increase in activation [40]. A similar pattern was also found in our own results across the imaging modalities. This highlights that the response to TUS is not uniform and may depend on individual-specific factors.

### Using Neuromodulation to Normalize Hyperactive Brain Circuits

While this study targeted the motor cortex in healthy individuals, its primary implication lies in demonstrating that tb-TUS can non-invasively *reduce* local brain connectivity and perfusion. This is particularly relevant for neuropsychiatric and neurological disorders characterized by network hyperactivity. For instance, evidence in Major Depressive Disorder (MDD) points to a frontostriatal salience network extension [49], and a protocol that reduces this pattern could be investigated to may normalize this extension. Similarly, Schizophrenia is associated with striatal hyperconnectivity and hyperconnectivity between the basal ganglia/thalamus and the sensor-motor network that can be indexed by neuroimaging [41–43], making the precise down-regulation of these circuits a potential non-pharmacological approach. The principle also extends to neurodegenerative conditions like Alzheimer’s Disease or Glioblastome, where amyloid-β (Aβ) or pathological neuron-glioma interactions induces a toxic hyperconnectivity that facilitates the spread of tau pathology [44] or the spread and recurrence of glioblastomas [45,46]. Attenuating this process is a key therapeutic goal that could be explored with inhibitory TUS. In another domain, the subjective experience of chronic pain is directly correlated with increased activity in cortical networks [47,48], suggesting that a tool to safely down-regulate these circuits e.g. in low-back pain patients could be explored as a non-opioid treatment.

### Limitations and Future Directions

The findings of this pilot study must be considered preliminary, primarily due to the modest sample size, which may have limited the statistical power to detect neurochemical changes. The immediate next step is to conduct larger-scale studies to replicate these primary CBF and FC findings. Future research must also prioritize multi-modal investigations that concurrently measure hemodynamic (fMRI/ASL) and electrophysiological (EEG/MEP) responses to clarify the complex effects of TUS. A systematic exploration of the TUS parameter space (e.g., intensity, frequency, duration) is essential to determine how to reliably produce specific inhibitory or excitatory effects.

It is crucial to emphasize that while the “theta-burst” terminology is borrowed from a similar pattern used in Transcranial Magnetic Stimulation (TMS) [49,50], tb-TUS and tb-TMS are fundamentally different technologies that operate on distinct physical principles to achieve neuromodulation. Therefore, our investigation is aimed at the specific, reported effects of the ultrasound protocol, independent of the rather well-documented outcomes of its TMS counterpart we also investigated recently [51,52]. Further, we conducted our study in late 2023 and adhered to a rather conservative intensity limit below 720 mW/cm² based on safety guidelines prevalent at the time. This ensures a non-significant thermal rise at the focus but may be considered a limitation as subsequent safety frameworks now permit less restrictive parameters [53]. Another potential limitation is the influence of auditory confounds [54], even so auditory confounds are more likely to occur at higher pulse repetition frequencies (PRFs) in the kilohertz (kHz) range.

## Conclusion

In conclusion, this pilot investigation demonstrates that tb-TUS is capable of producing quantifiable, modulatory effects on hemodynamic and connectivity measures in the human brain. While preliminary, these findings encourage further investigation. A crucial next step is to identify reliable neuroimaging markers of TUS effects. This is essential for developing TUS from a promising neuromodulation technique into a targeted, circuit-based therapy for various brain disorders.

## Acknowledgments

The study was endorsed by the Federal Ministry of Education and Research (Bundesministerium für Bildung und Forschung [BMBF]) within the initial phase of the German Center for Mental Health (DZPG) (grant: 01EE2303A, 01EE2303F to PF, AS).

## Supporting Information

**Table S1:**
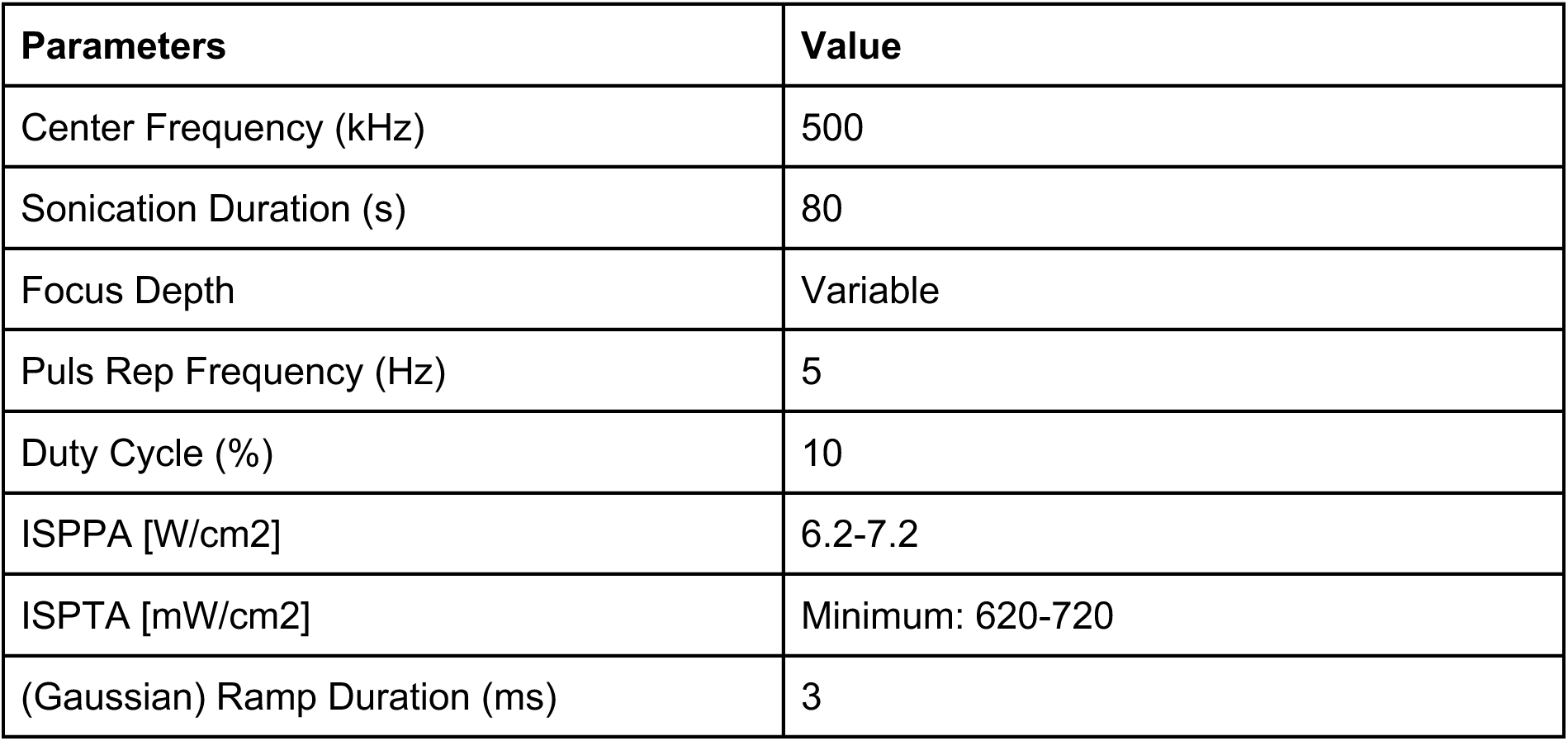
Tb-TUS Stimulation Parameters. Stimulation parameters of both sham and active tb-TUS stimulation. ISPPA = spatial Peak Pulse Average Intensity, ISPTA = Spatial Peak Temporal Average Intensity.

| Parameters | Value |
| --- | --- |
| Center Frequency (kHz) | 500 |
| Sonication Duration (s) | 80 |
| Focus Depth | Variable |
| Puls Rep Frequency (Hz) | 5 |
| Duty Cycle (%) | 10 |
| ISPPA [W/cm<sup>2</sup>] | 6.2-7.2 |
| ISPTA [mW/cm<sup>2</sup>] | Minimum: 620-720 |
| (Gaussian) Ramp Duration (ms) | 3 |

**Table S2.** MRI Acquisition Parameters. Scanning parameters used for structural images (T1w, T2w), pcASL, rs-fMRI and MRS. pcASL = pseudo-continous Arterial Spin Labeling. MRS = Magnetic Resonance Spectroscopy. EPI = Echo Planar Imaging. FT = Fingertapping task. *Note: Field maps were additionally recording at the beginning, during and after all scans. Both structural images, the T1-weighted and T2-weighted recordings, were acquired to create the pseudo-CT scans and for neuroradiologists to identify possible microstructural injuries of the brain after sonication. For safety evaluation the three additional sequences were recorded before and after active and sham tb-TUS: DWI-difussion and T2-SWI. DWI= Diffusion-Weighted Imaging. SWI= Susceptibility Weighted Imaging*.

|  | T1weighted | T2weighted | pcASL | EPI (rs-fMRI<br>AP & PA) | MRS<br>Spectroscopy | FT |
| --- | --- | --- | --- | --- | --- | --- |
|  | MP-RAGE | FLAIR | CMRR<br>Multiband<br>EPI | CMRR<br>Multiband<br>EPI | CMRR msLaser | Single Band<br>EPI |
| TR [ms] | 2300 | 5000 | 8000 | 800 | 3000 | 3000 |
| TE [ms] | 2.26 | 388 | 54 | 37 | 68 | 30 |
| Flip angle | 8° | 120° | 90° | 52 | 90/180 | 90° |
| Resolution [mm<sup>3</sup>] | 1 x 1 x 1 | 1 x 1 x 1 | 2.5x2.5x2.3 | 2 x 2 x 2 | 20 x 30 x 20 | 2 x 2 x 2 |
| Thickness [mm] | 1.0 | 1.0 | 2.27 | 2.0 | - | 2 |
| Accelerator/multiband factor | 2/- | 2/- | 6 / - | -/8 | - /- | 2/-2 |
| PAT mode | GRAPPA | GRAPPA | - | - | - | GRAPPA |
| Field of View [mm] | 256 | 256 | 215 | 208 | - | 200 |
| Number<br>of slices /averages | 192 | 176 | 60 | 208 | 8 / 128 | 72 |
| Imaging Direction | Sagittal | Sagittal | Axial | Axial | Axial | Transversal |
| Duration | 5:21 | 05:22 | 05:29 | 6:40/<br>6:40 | 9:33 | 5:56 |
| Measurement(s) | 1 | 1 | 1 | 488/488 | 1 | 115 |
| Total Time | 45:01 |  |  |  |  |  |

**Table S3.**
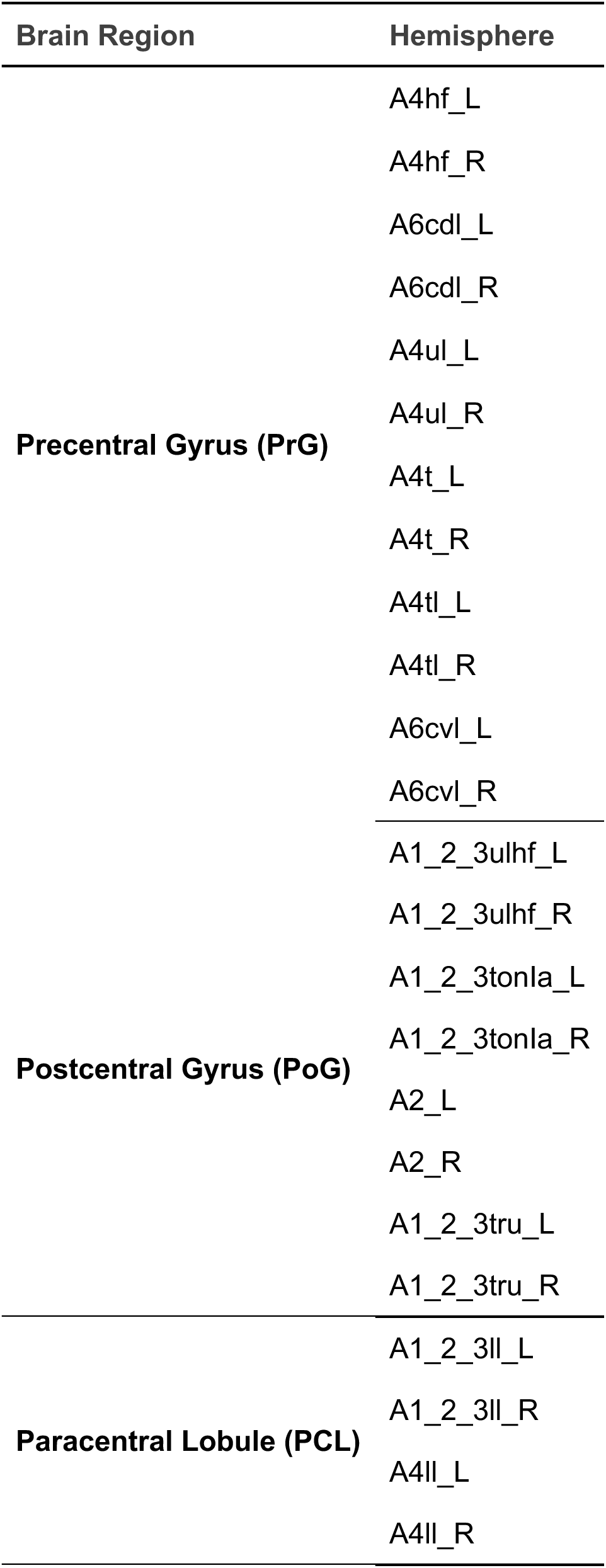
Brain Regions in the Primary Motor Cortex Used for Further Analysis for ASL and fMRI. Adapted regions focus on the PrG, PoG and PCL. Regions were adapted from the Brainetome atlas [27].

**Table S4.** Tb-TUS studies using similar protocols as our current study. This table provides an overview of studies in the literature using comparable stimulation parameters. The first author, journal and year of publication is provided, alongside with the anatomical target of the tb-TUS application, key sonication parameters, the primary outcome measures used, and the principal findings from each study.

| Study (Author, Journal, Year) | FUS Target | Key FUS Parameters | Outcome Measures | Key Findings (Direction of Change) |
| --- | --- | --- | --- | --- |
| Current Study | Left M1 | **500 kHz, 80s duration, Continuous tb-TUS (5 Hz PRF, 10% DC)**, ISPTA 620-720 mW/cm<sup>2</sup> | pcASL-CBF, rs-fMRI FC, MRS | ↓ CBF, ↓ Within-Region FC, ~ (non-sig ↓) GABA |
| Zeng et al., *Annals of Neurology* 2022 [19] | Left M1 | **Similar Continuous tb-TUS protocol** | MEPs | ↑ MEP amplitude |
| Samuel et al., *Brain Stimulation* 2022 [20] | Left M1 | **Similar tb-TUS parameters** | MEPs, MEG | ↑ MEP amplitude, ↓ Alpha/Beta band desynchronization |
| Shamli Oghli et al., *Brain Stimulation* 2023 [55] | Left M1 | **500 kHz, 80s Continuous tbTUS (5 Hz PRF, 10% DC)**, in situ ISPPA 2.93 W/cm<sup>2</sup> | MEPs (pharmaco-TMS study) | Placebo: ↑ MEP amplitude; Effect blocked by Na<sup>+</sup>, Ca<sup>2+</sup>, GABA, & NMDA drugs |
| Yaakub et al., *Nature Communications* 2023 [21] | PCC / dACC | **Similar 80s tb-TUS protocol** | MRS (GABA), rs-fMRI FC | ↓ GABA (in PCC only), ↑ FC (in both regions) |
| Bao et al., *The Journal of Physiology* 2024 [56] | Left M1 (superficial & deep layers) | **500 kHz, Continuous tb-TUS (5 Hz PRF, 10% DC)**; Intermittent tb-TUS (5 Hz PRF, 10% DC) | MEPs | Continuous: ↓ MEP amplitude; Intermittent: No change |
| Xia et al., *The Journal of Physiology* 2024 [57] | Left M1 | **500 kHz, 80s tbTUS (5Hz PRF, 10% DC)**, in situ ISPPA 2.93 W/cm<sup>2</sup> | MEPs (bilateral), IHI, ICF | ↑ Left M1 excitability (local); ↓ Right M1 excitability (remote) |
| Zeng et al., *Brain Stimulation* 2024 [58] | Left M1 | Parametric study of tbTUS: Varied Intensity (10 vs 20 W/cm<sup>2</sup>), PRF (2, 5, 10 Hz), Duty Cycle (5, 10, 15%), & Duration (40, 80, 120s) | MEPs, SICl, ICF, SICF | Dose-dependent ↑ increase in MEPs; 5 Hz PRF was optimal. Higher intensity, DC, & duration led to stronger/longer effects. |
| Cain et al., *Scientific Reports* 2021 [35] | Left Globus Pallidus | 650 kHz, 5% DC, ISPTA ~720 mW/cm<sup>2</sup>, 10 Hz or 100 Hz PRF | BOLD fMRI, ASL-perfusion | ↓ BOLD signal (100 Hz only), ↓ generalized perfusion (both) |
| Darmani et al., *Nature Communications* 2025 [[36] | Globus Pallidus Internus (GPi) | **500 kHz, tb-TUS (5 Hz PRF, 10% DC)** and 10Hz TUS (10Hz PRF, 30% DC) | Local Field Potentials (LFPs), Behavior (SSRT) | ↑ LFP power (tbTUS ↑ theta; 10Hz TUS ↑ beta), ↑ SSRT (impaired inhibition) |

**Figure S1.**
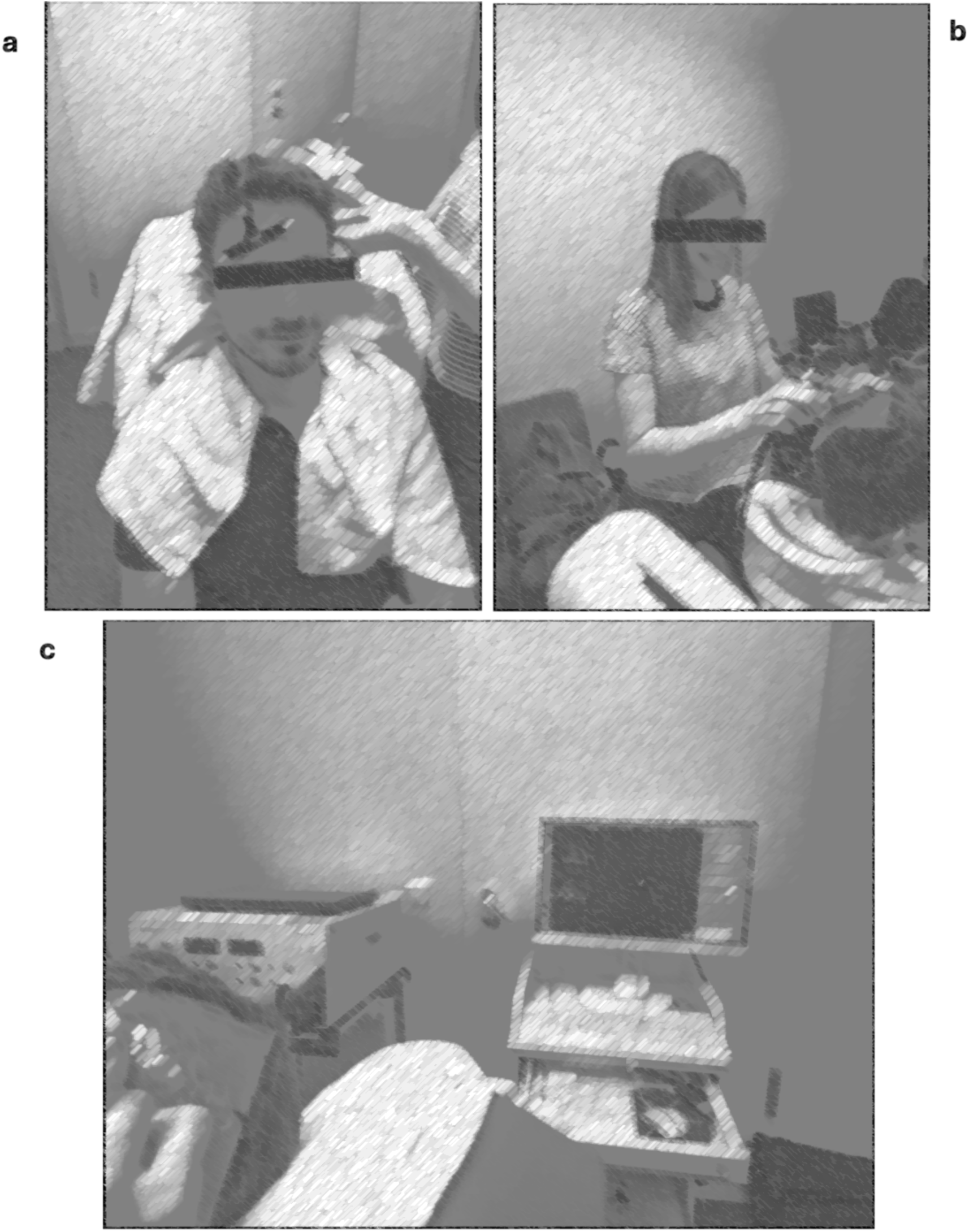
Set-Up of tb-TUS Stimulation Guided by Neuronavigation. **a.** The participant was placed in an upright position. **b.** The participant is equipped with gel pads to ensure a smooth transition, and the transducer is constantly kept in the position by the researcher. **c.** To ensure stimulation of the targeted area, the Tb-FUS stimulation is guided by neuronavigation using LOCALITE (LOCALITE, Bonn, Germany). *Nnote: the pictures have been made unidentifiable in order to protect the anonymity of our test subjects*.

**Figure S2.**
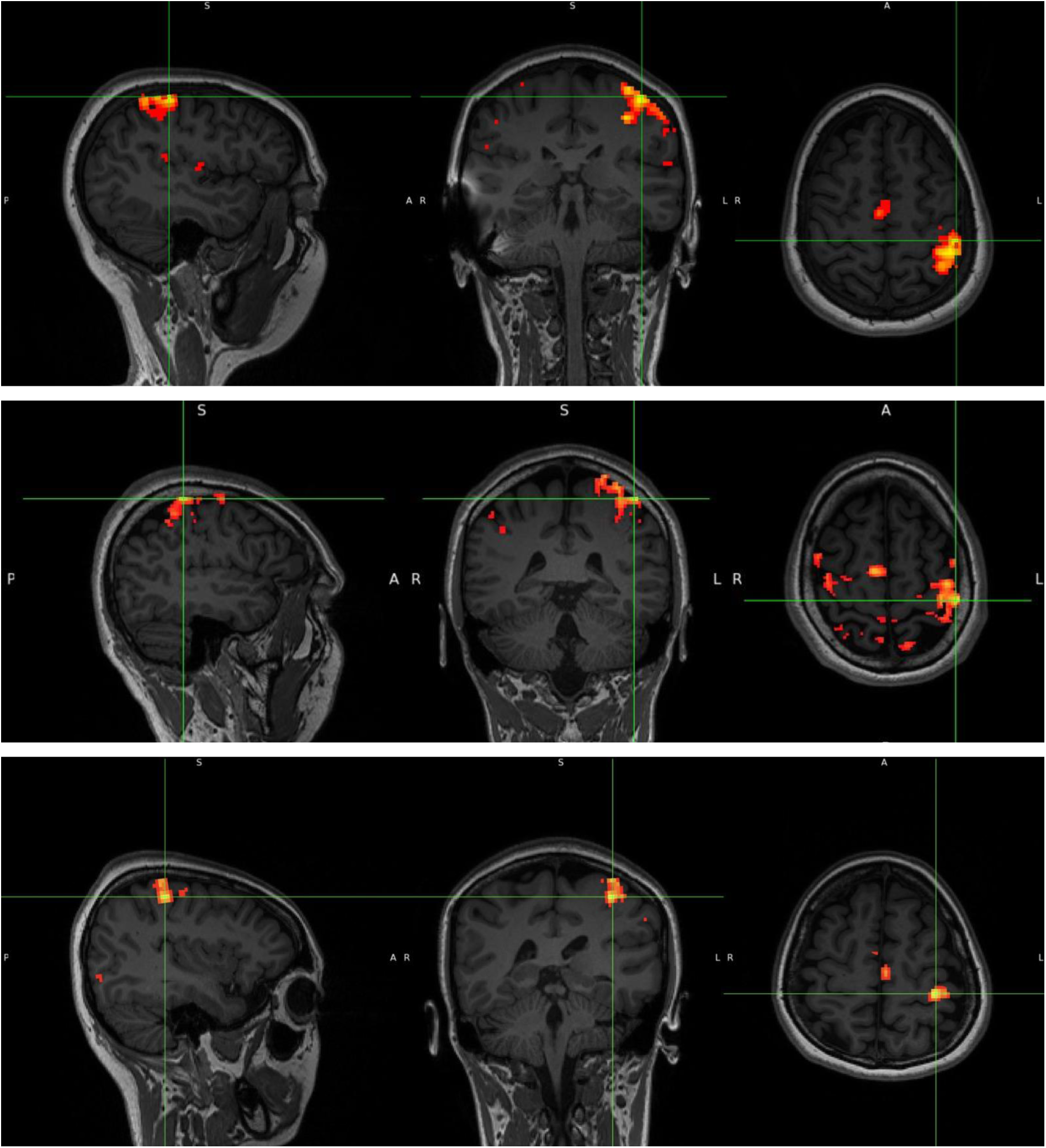
Examples of individual M1 targeting. Targeting for M1 was individually determined by the result of the finger tapping task and a thresholding of z>6. This peak region was used for the simulation and the subsequent neuronavigated stimulation. Above are three example results with the marked peak activations.

**Figure S3.**
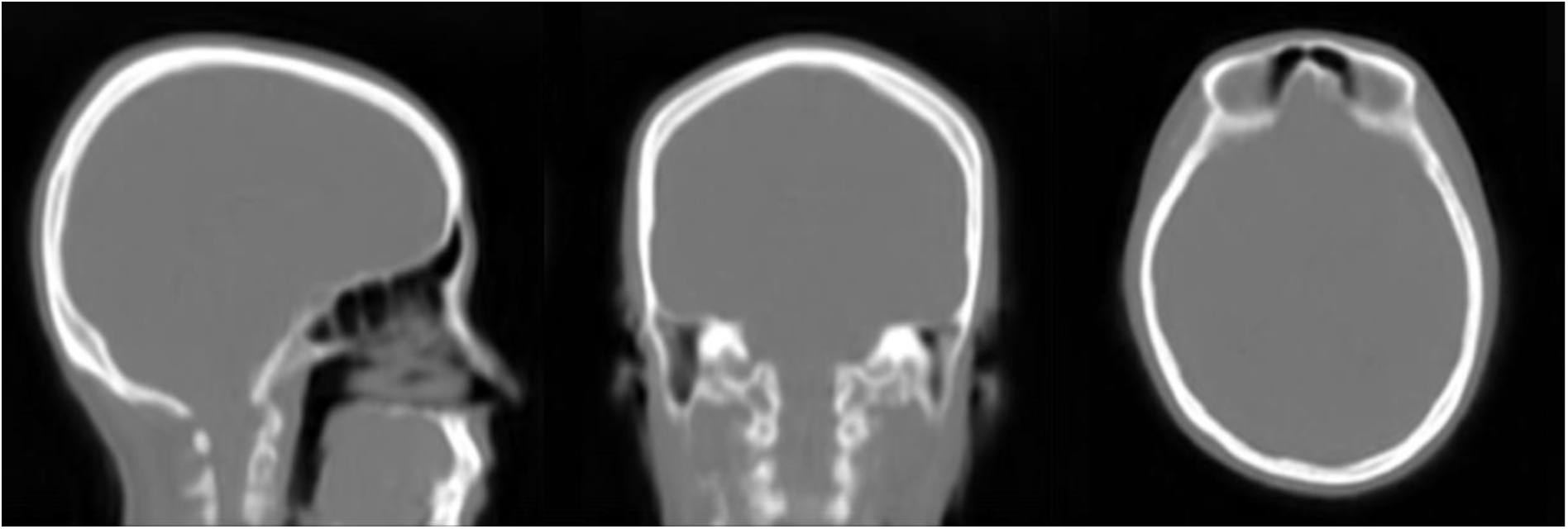
Representation of the Pseudo Computer Tomography (CT) Image. A Pseudo CT Image of a representative participant was created based on the individual T1-weighted images of each corresponding subject. Pseudo CTs were created using the MR-to-pCT for FUS acoustic simulations code (version 1.0.0), available on GitHub [24].

**Figure S4.**
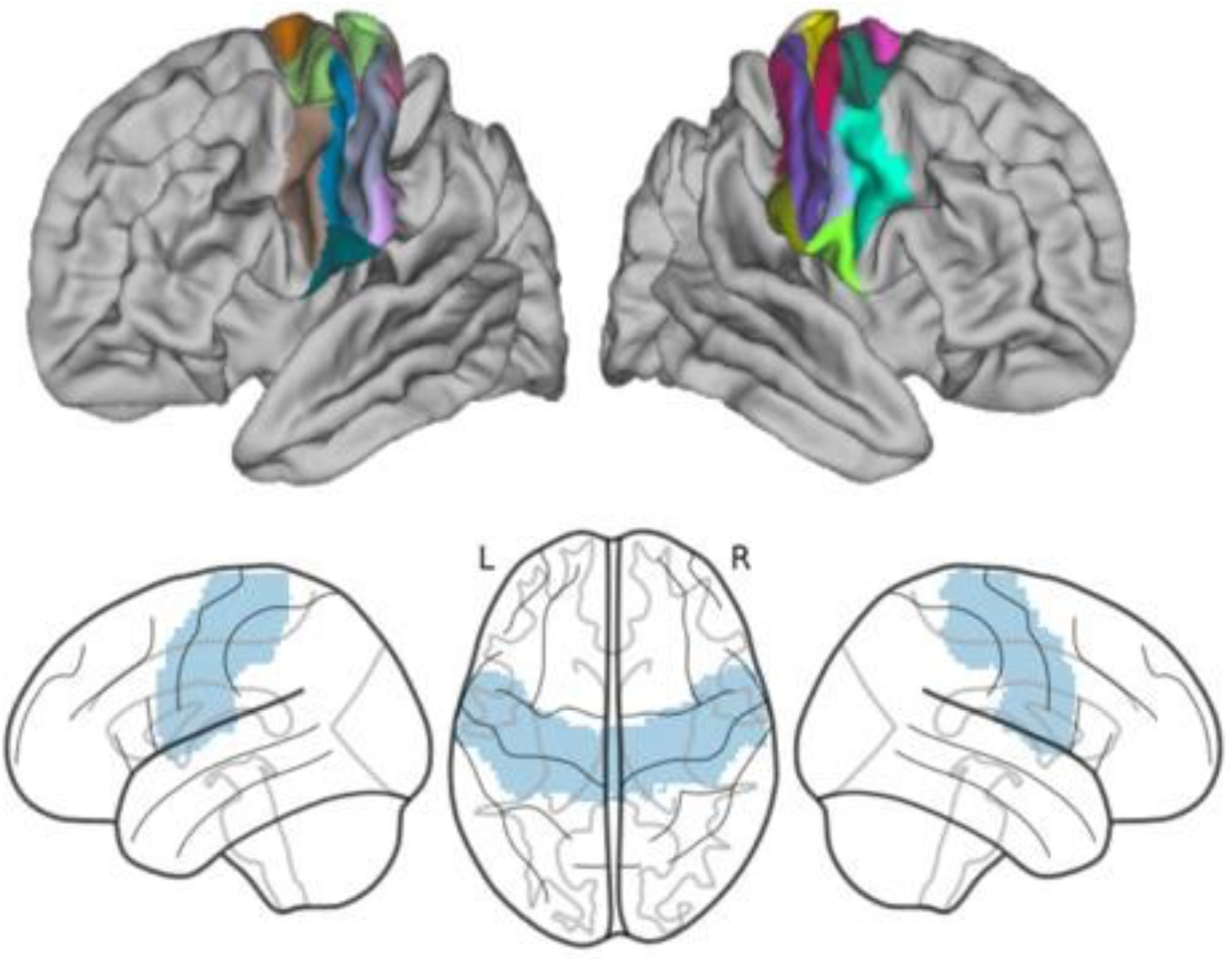
Regions of Interest in the Somatomotor Cortex. ROIs used for further analysis include the PrG, PoG, and PCL based on the standardized Brainetome Atlas [27]. The top panel represents the ROIs corresponding to the Brainetome regions (see also Table S3) and was created using *Connectome Workbench* (https://www.humanconnectome.org/software/connectome-workbench). The bottom panel depicts the combined ROIs visualized using glass brain plots created with *Nilearn* (http://nilearn.github.io).

